# Exploratory metagenomics of bacterial diversity in semen from indonesian native roosters supplemented with curcumin and penicillin-streptomycin

**DOI:** 10.64898/2026.08.27.747682

**Authors:** Khaeruddin, Hermawansyah, Junaedi, Bahri Syamsuryadi, Kasri

## Abstract

**Objective:** This study aims to evaluate the effectiveness of curcumin and penicillin-streptomycin as diluents on changes in the structure and diversity of the chicken semen microbiome during storage.

**Methods:** Semen was collected from Kampung chickens (native to Indonesia) and divided into five treatment groups: diluted without antibiotics or curcumin (control), and diluted with additions of 10 μM, 20 μM, and 30 μM curcumin, and penicillin-streptomycin, respectively. The semen was stored at 5 °C for 24 hours. The composition and diversity of the semen microbiome were analyzed using 16S full-length amplicon sequencing.

**Results:** Analysis of the top 10 species showed that Uncultured Saccharofermentans sp. and Porphyromonas somerae served as the most dominant and stable core microbiome across all treatments. Alpha diversity analysis showed that the addition of curcumin and penicillin-streptomycin reduced microbial richness (Observed, Chao1, ACE, and Fisher) in a dose-dependent manner, yet maintained overall diversity (Shannon and Simpson), with the penicillin-streptomycin treatment resulting in the highest species evenness (InvSimpson). Beta diversity analysis revealed extreme separation of taxonomic abundance variance in the penicillin-streptomycin group, whereas the curcumin treatment exhibited a dose-dependent pattern of microbial abundance transition. Venn diagram analysis identified 415 OTUs as the core microbiome and confirmed that curcumin acts through selective filtering that stabilizes the ecosystem without triggering the proliferation of opportunistic taxa.

**Conclusion:** Penicillin-streptomycin acts more rapidly and dominantly in suppressing/killing bacterial populations; however, the addition of curcumin is able to modulate the microbial ecosystem in a more balanced manner by suppressing the growth of harmful bacteria without compromising the integrity of the chicken semen environment.

## INTRODUCTION

The dilution of chicken semen is necessary to optimize semen quality during short- and long-term storage to support the success of artificial insemination (AI). The semen collection process can be contaminated by microorganisms, such as bacteria originating from feces, because the chicken’s fecal and semen excretory tracts are connected. These bacteria can multiply during semen storage prior to AI. Bacterial contamination is common in poultry semen and can reduce sperm quality and fertility [1]. Although storage temperatures are lowered to inactivate sperm during storage, bacterial growth can still occur [2]. During storage, the quality of chicken semen is compromised by endotoxins produced by bacteria [3].

The impact of bacterial contamination on semen quality is a rapidly evolving field of research [4]. Studies in humans have revealed that a diverse microbial community in semen can interact with spermatozoa and affect their function [5]. A progressive decrease in chicken sperm motility (45%) was found to correlate with an increase in bacterial colony count (population of 13.35 log10 CFU/ml) [6]. Media containing high bacterial counts (10 CFU/ml) have also been reported to cause a decrease in chicken sperm motility [7]. Bacteria in semen can trigger biological stress and affect semen quality parameters, including sperm motility, mitochondrial activity, membrane and acrosome integrity, and DNA fragmentation [6].

One strategic measure to address bacterial contamination during semen collection and storage is the addition of antibiotics to the semen diluent [8]. Antibiotics help prevent bacterial contamination and protect females from bacterial pathogenesis [1]. Penicillin and streptomycin are the types of antibiotics frequently added to the semen diluents of indigenous Indonesian chickens during the storage process [9]; however, their effects on the abundance of the microbiome in the seminal plasma of these chickens have not yet been studied. Another challenge is that the use of non-therapeutic antibiotics at low doses can still lead to microbial resistance [10].

Most available antibiotics have led to the development of resistance [11], creating a need for natural biomolecules as alternatives to antibiotics that possess antibacterial and antioxidant properties [8]. Plant-based compounds can serve as alternatives to commercial antibiotics because they possess antimicrobial properties while also preventing oxidative stress by enhancing the activity of several antioxidant enzymes [12]. One plant compound with potential to replace antibiotics is curcumin, as it has been proven to possess broad-spectrum antibacterial properties [13]. Curcumin supplementation reduces the growth of aerobic bacteria that can be cultured during refrigerated storage, with more pronounced effects observed after longer storage periods [14].

Metagenomic studies of semen have been conducted on commercial chicken strains to examine microbial resistance to the antibiotics ampicillin, chloramphenicol, gentamicin, imipenem, levofloxacin, tetracycline, tigecycline, and tobramycin [20]. Although previous research has confirmed the protective role of curcumin against chicken sperm damage caused by three species of pathogenic bacteria in vitro using isolation methods [15], there remains a lack of comprehensive information regarding these bacteria. The main novelty of this study lies in the use of NGS-based metagenomics to reveal how increasing doses of curcumin and penicillin-streptomycin alter the entire profile of the chicken semen bacterial community independently of culture and to comprehensively characterize bacterial populations using nanopore sequencing. This study aims to evaluate the effectiveness of curcumin and penicillin-streptomycin as diluents in altering the structure and diversity of the chicken semen microbiome during storage using a metagenomic approach.

## MATERIALS AND METHODS

### Chicken Rearing and Semen Collection

19 Kampung roosters (a native Indonesian breed) aged approximately 10 months were reared in cages measuring 50 x 60 x 60 cm³. They were fed a complete feed at a rate of 150 g per bird per day (crude protein 16%) and provided with water ad libitum. Semen was collected using the cloacal massage technique [16]. The semen was collected using a 1-ml syringe and combined into a single tube.

### Dilution and Cooling

The diluent used was Ringer’s lactate (Widatra Bakti, Indonesia) (8.6% NaCl, 0.31% NaC H O, 0.03% KCl, and 0.02% CaCl in sterile water); the pH was adjusted to 7.9 by adding tris hydroxyl aminomethane. The collected semen was divided into 5 tubes, each treated as follows: control, 10 μM curcumin (Sigma-Aldrich, USA), 20 μM curcumin, 30 μM curcumin, and penicillin-streptomycin. The penicillin (Meiji, Indonesia) dose was 1000 IU/ml and the streptomycin (Meiji, Indonesia) dose was 1 mg/ml [9] (Khaeruddin et al., 2024). Next, the liquid semen was stored at 5 °C for 24 hours. Each semen sample was mixed with DNA/RNA Shield (Zymo Research, US) in a 1:3 ratio.

### DNA Extraction and Sequencing

DNA was extracted from liquid semen samples using the Quick-DNA Magbead Plus Kit (Zymo Research, D4082, US), and its concentration was determined using a NanoDrop spectrophotometer and a Qubit fluorometer. Only DNA concentrations of at least 2 ng/μL (measured with the NanoDrop) or 0.1 ng/μL (measured with the Qubit) were used for the PCR amplification step. The highly variable V1–V9 regions of the bacterial 16S rRNA gene were amplified using the primer pair 27F (5’-AGAGTTTGATCMTGGCTCAG-3’) and 1492R (5’-GGTTACCTTGTTACGACTT-3’) [17], using Phusion Plus PCR Master Mix (ThermoScientific, F632L, US). The quality and size of the PCR products were evaluated using agarose gel electrophoresis. A total of 2 μL of each PCR product was loaded onto a 1% agarose gel in 1× TBE buffer. A 1-kb DNA ladder (1 μL) was used as a molecular weight marker. Electrophoresis results showing PCR product bands in the 1500–1600 bp range could proceed to the next stage. Accurate quantification and purification of PCR products were performed using Qubit; only purified amplicons meeting the minimum requirement of 20 ng/μL in a final volume of 10 μL were used for library preparation. Library preparation was performed using the Nanopore Sequencing Ligation—Native Barcoding Kit 96 V14 (Oxford Nanopore Technologies, SQK-NBD114.96, UK), and the final library was sequenced using the Nanopore GridIon platform. Nanopore sequencing is run using MinKNOW software version 26.01.15. Base calling is performed using Dorado version 7.13.6 with a high-accuracy model.

### Bioinformatics processing

For bioinformatics analysis, adapters and primers were trimmed using Dorado version 7.13.6, and reads were filtered based on quality and length using Nanofilt and visualized using NanoPlot. The filtered reads were grouped using the Proname version 2.3.0 workflow and classified against the rEGEN-B database with a minimum query coverage and a 90% identity percentage. Further analysis and visualization were performed using Krona Tools and RStudio with R version 4.3. Alpha diversity indices, including Chao1 (community richness), as well as Shannon and Simpson (community diversity and evenness), were calculated to evaluate microbial diversity in the samples. Beta diversity was analyzed to measure variations in microbial community structure across sample groups. Beta diversity measures biological diversity across regions or ecosystems. The beta diversity analysis consisted of three parts: PCA, PCoA, and UPGMA.

## RESULTS

### Venn Diagram

Based on the two-sided Venn diagram (Figure 1) comparing the control and penicillin-streptomycin groups, a total of 739 core taxa were identified. The highest diversity of unique taxa was found in the control group (471 taxa) and the lowest in the penicillin-streptomycin group (336 taxa). Overall, a total of 415 core taxa were identified—that is, microbial communities that were consistently present in all treatment groups. The highest diversity of unique taxa was found in the 30 μM curcumin group (59 taxa), followed by the 10 μM curcumin group (88 taxa), the 20 μM curcumin group (98 taxa), the penicillin-streptomycin group (109 taxa), and the control group (117 taxa).

**Figure 1.**
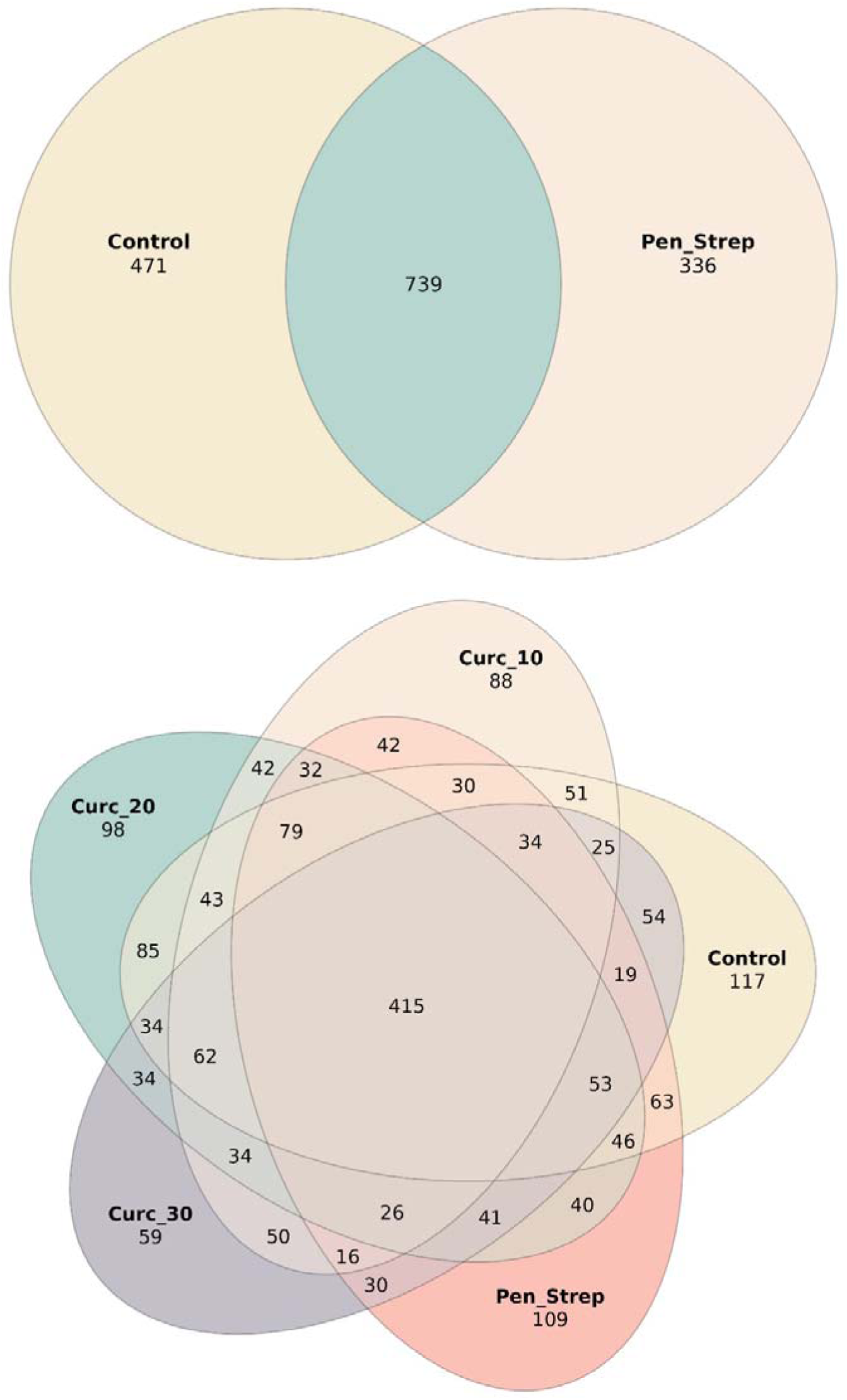
Two-sided venn diagram of metagenomic taxon distribution (top), 5-sided venn diagram of metagenomic taxon distribution (bottom).

### Krona Plot

Visualization of the taxonomic hierarchy using a Krona diagram (Figure 2) shows that in the control group, the Krona diagram was dominated by Phylum Bacillota (69.96%), Class Clostridia (30.26%), Order Lactobacillales (18.63%), Family Lactobacillaceae (17.86%), and Genus Ligilactobacillus (14.13%). Treatment with curcumin and penicillin-streptomycin caused an expansion of the area occupied by the Order Eubacteriales (18.66–20.90%) and the Family Oscillospiraceae (11.45–12.33%), making them more dominant. At the genus level, the control group was dominated by Ligilactobacillus. At curcumin concentrations of 10–20 μM, the expansion was dominated by Porphyromonas (12.25–12.33%); however, at a curcumin concentration of 30 μM and with the addition of penicillin-streptomycin, the expansion was dominated by Saccharofermentans (11.45–12.10%).

**Figure 2.**
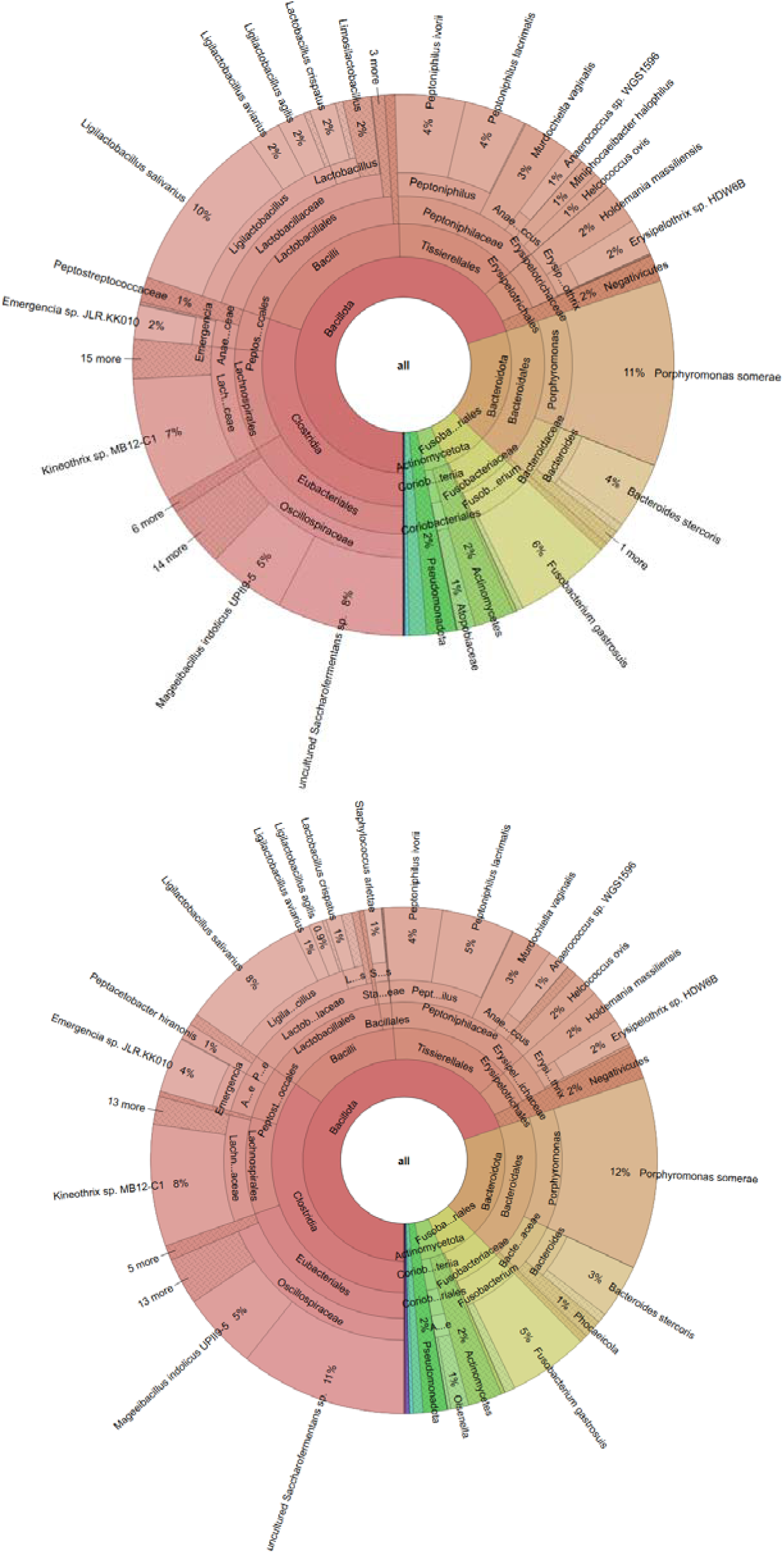
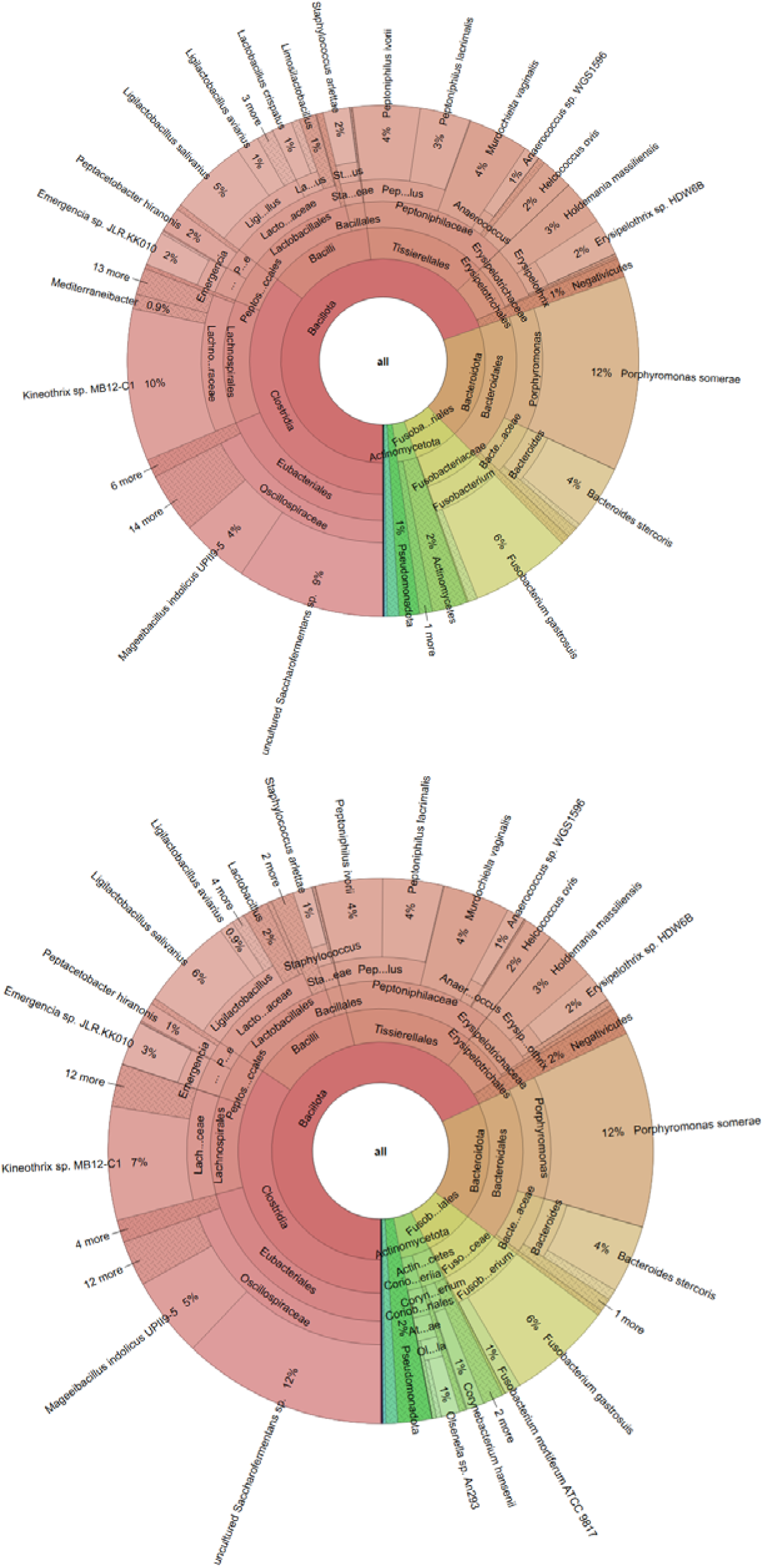
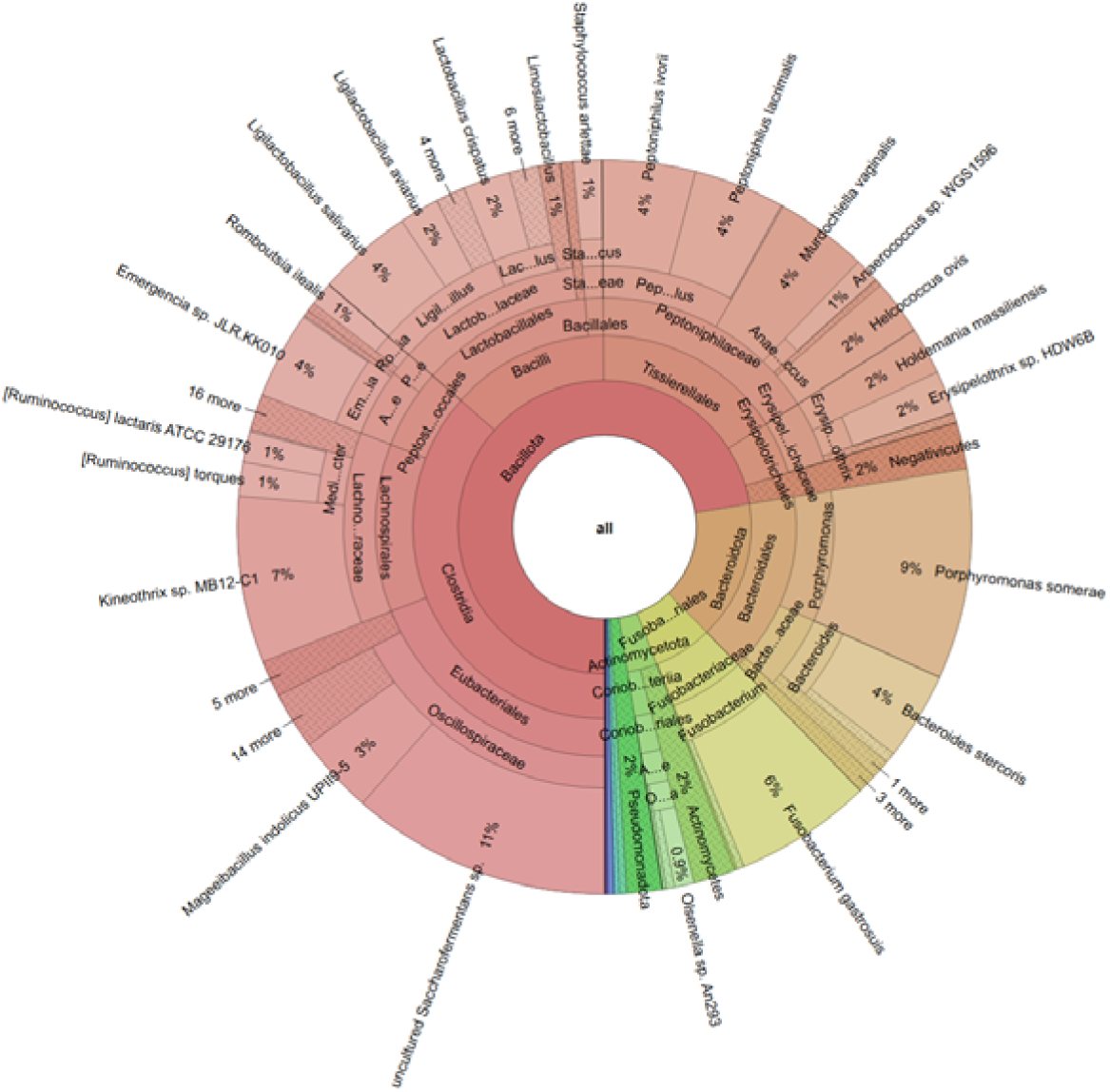
Krona plots of bacteria in rooster semen. From top to bottom, respectively: control, curcumin 10 μM, 20 μM, 30 μM, and penicillin-streptomycin.

### Sankey Diagram

The Sankey diagram (Figure 3) shows that the bacterial community in the control group’s chicken semen was dominated by the phylum Bacillota, with the main contribution coming from the class Bacilli, particularly the family Lactobacillaceae and the genus Ligilactobacillus. At the species level, Ligilactobacillus salivarius was one of the most dominant taxa. In addition to Bacillota, the community also consisted largely of Bacteroidota, particularly the family Porphyromonadaceae and the genus Porphyromonas, as well as Fusobacteriota, which was primarily represented by Fusobacterium. Meanwhile, Actinomycetota, Campylobacterota, and Pseudomonadota were found in lower abundances.

**Figure 3.**
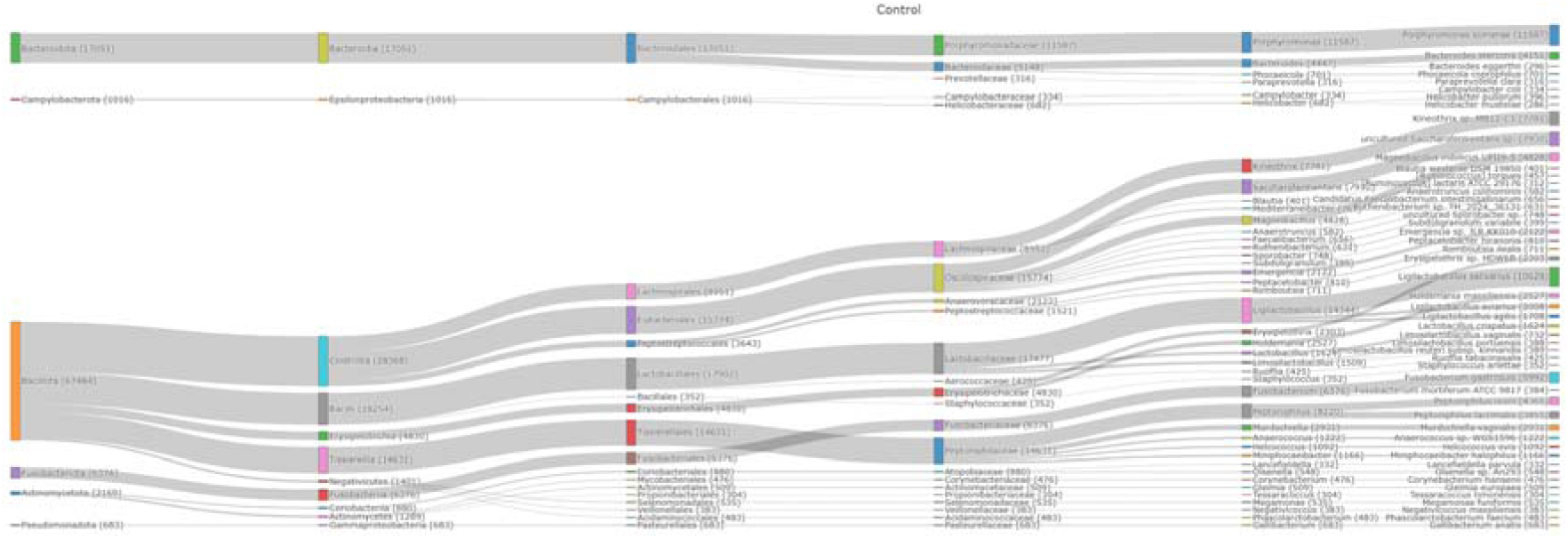
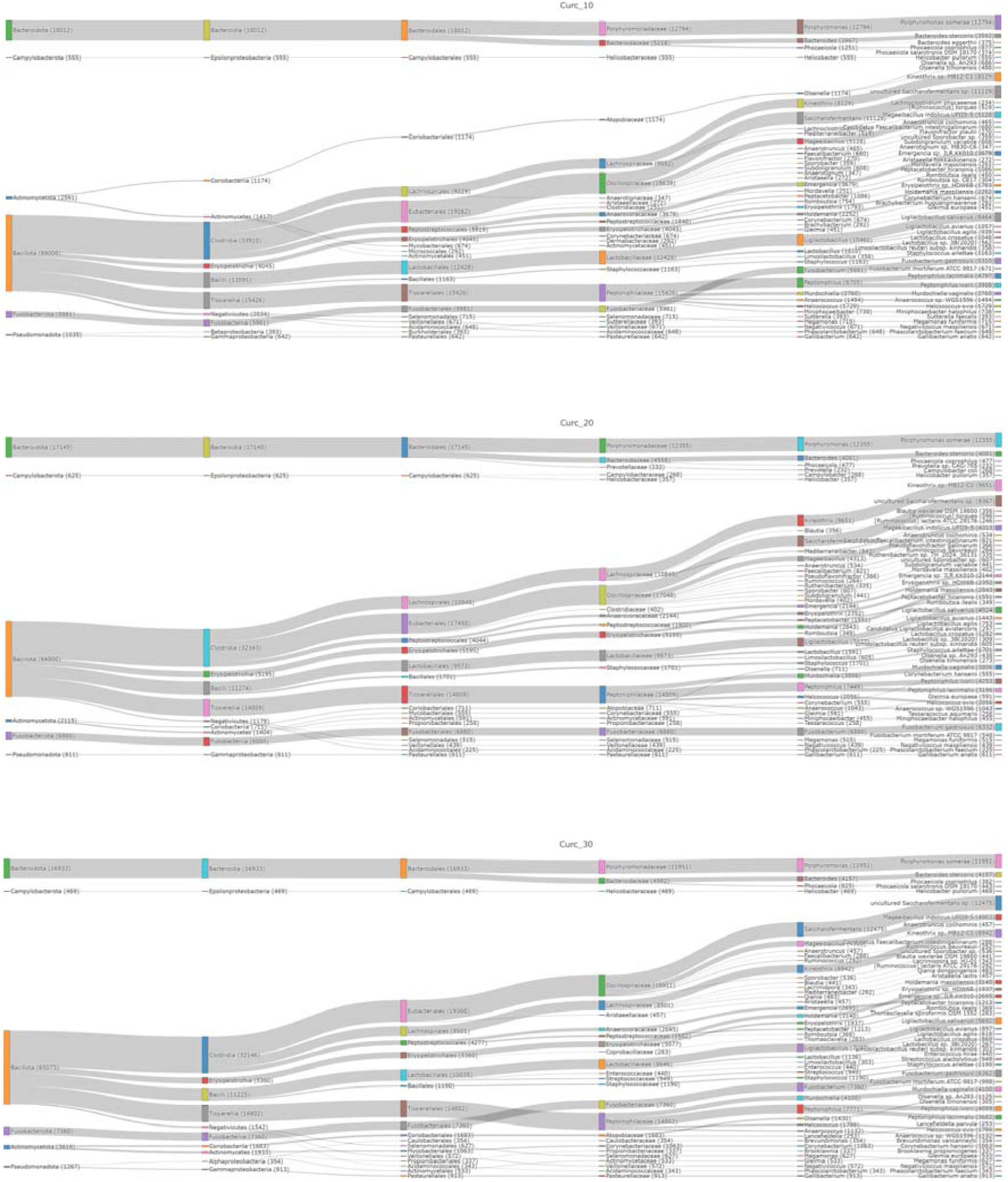
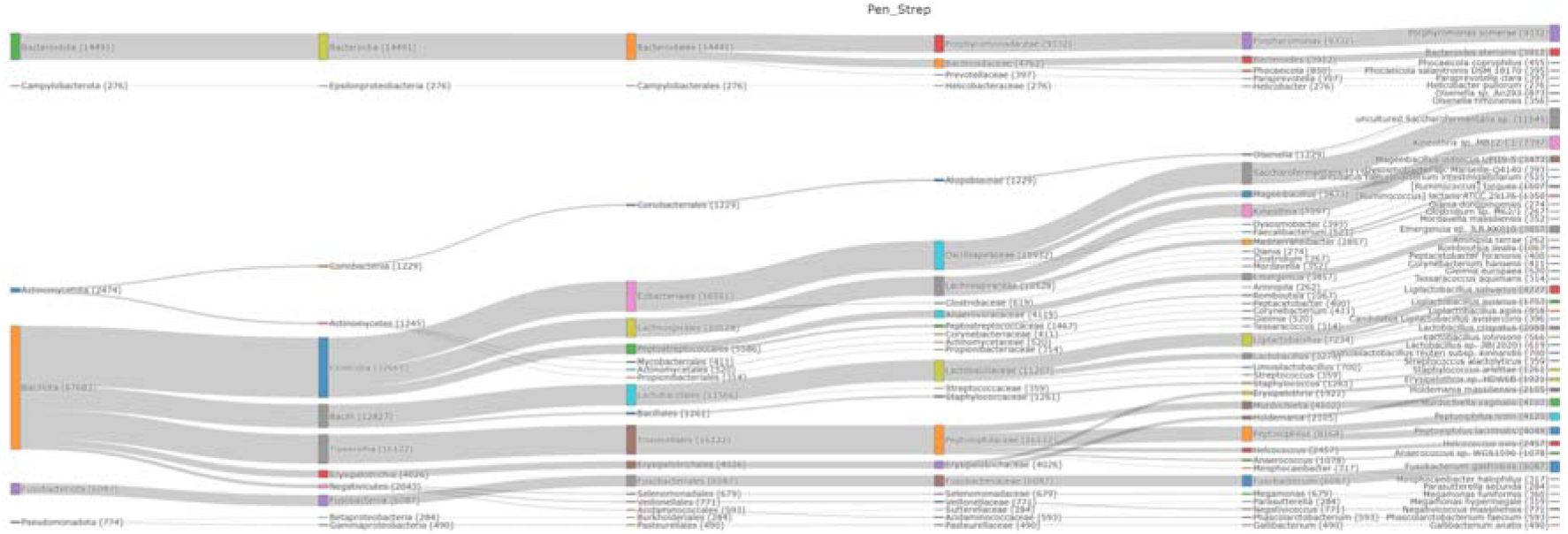
Sankey diagram of bacteria in rooster semen. From top to bottom: Control, penicillin-streptomycin, curcumin 10 μM, 20 μM, and 30 μM.

### Top 10 Taxa

Metagenomic analysis of the top 10 bacterial species (Figure 4) showed that Uncultured Saccharofermentans sp. and Porphyromonas somerae served as the most dominant and stable core microbiome across all treatments. Porphyromonas somerae remained relatively stable (11.10– 12.25%) in the control group and in treatments with increasing curcumin concentrations. In contrast, with the use of penicillin-streptomycin, its abundance decreased slightly (9.26%), indicating that the addition of antibiotics tended to suppress the proportion of Porphyromonas somerae. The lowest proportion of uncultured Saccharofermentans sp. was observed in the control group (7.60%). Administration of curcumin and penicillin-streptomycin consistently increased its abundance (≥ 9.34%). Ligilactobacillus salivarius showed a tendency toward a decrease in relative abundance due to treatment (≤ 8.10%) compared to the control (10.18%). The most significant decrease was observed with penicillin-streptomycin and 20 μM curcumin (∼4%). This pattern indicates that these treatment conditions exerted selective pressure that inhibited the growth of the Ligilactobacillus salivarius population.

**Figure 4.**
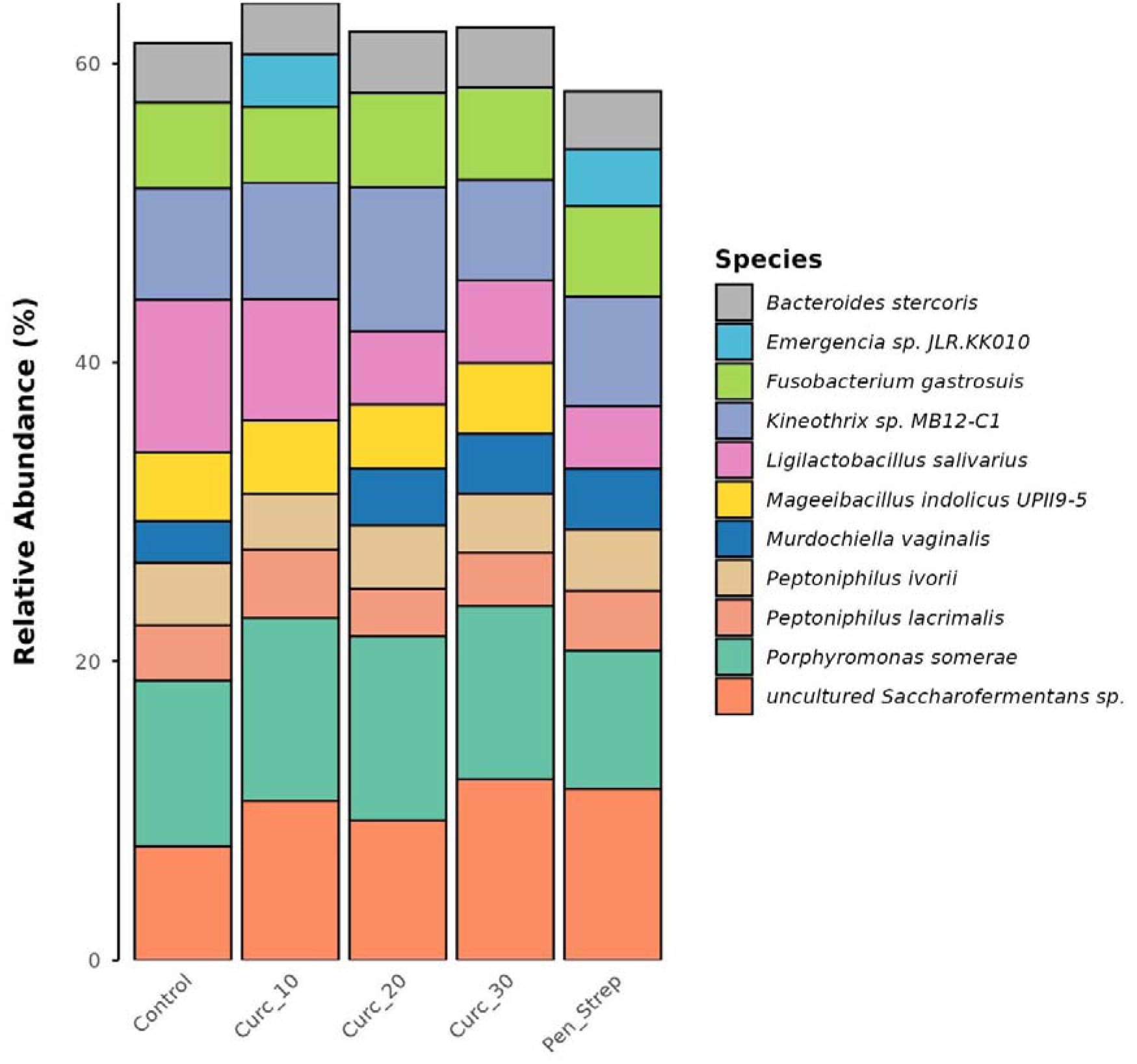
Relative abundance of the top 10 bacteria in rooster semen

**Figure 5.**
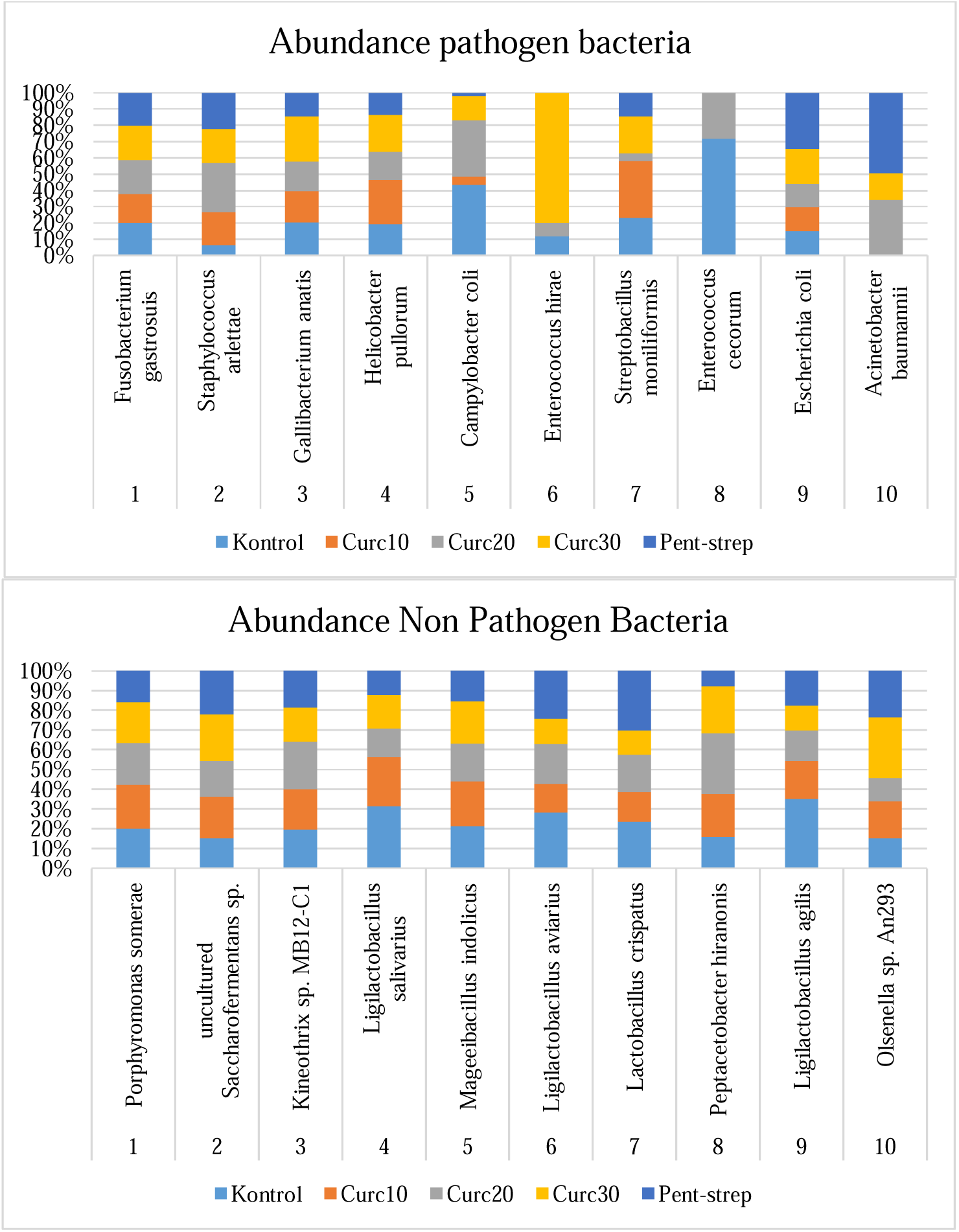
Relative abundance of the top 10 pathogenic bacteria (top) and nonpathogenic bacteria (bottom) in rooster semen.

Based on the bacterial community profile, the pathogen/pathobiont group (Table 1) showed varying responses to the curcumin dose treatments. Fusobacterium gastrosuis was one of the most dominant and relatively stable taxa across all treatments, with its abundance increasing at 20 μM and 30 μM curcumin compared to the control. Gallibacterium anatis also showed an increase at a 30 μM curcumin dose, while Campylobacter coli experienced the most significant decrease at 10 μM curcumin but increased again at 20–30 μM. Helicobacter pullorum increased at 10 μM curcumin, then decreased at 20 μM curcumin before rising slightly again at 30 μM curcumin. Meanwhile, Enterococcus hirae was not detected at 10 μM curcumin but increased sharply at 30 μM curcumin. Escherichia coli remained relatively stable across the curcumin groups.

**Table 1.**
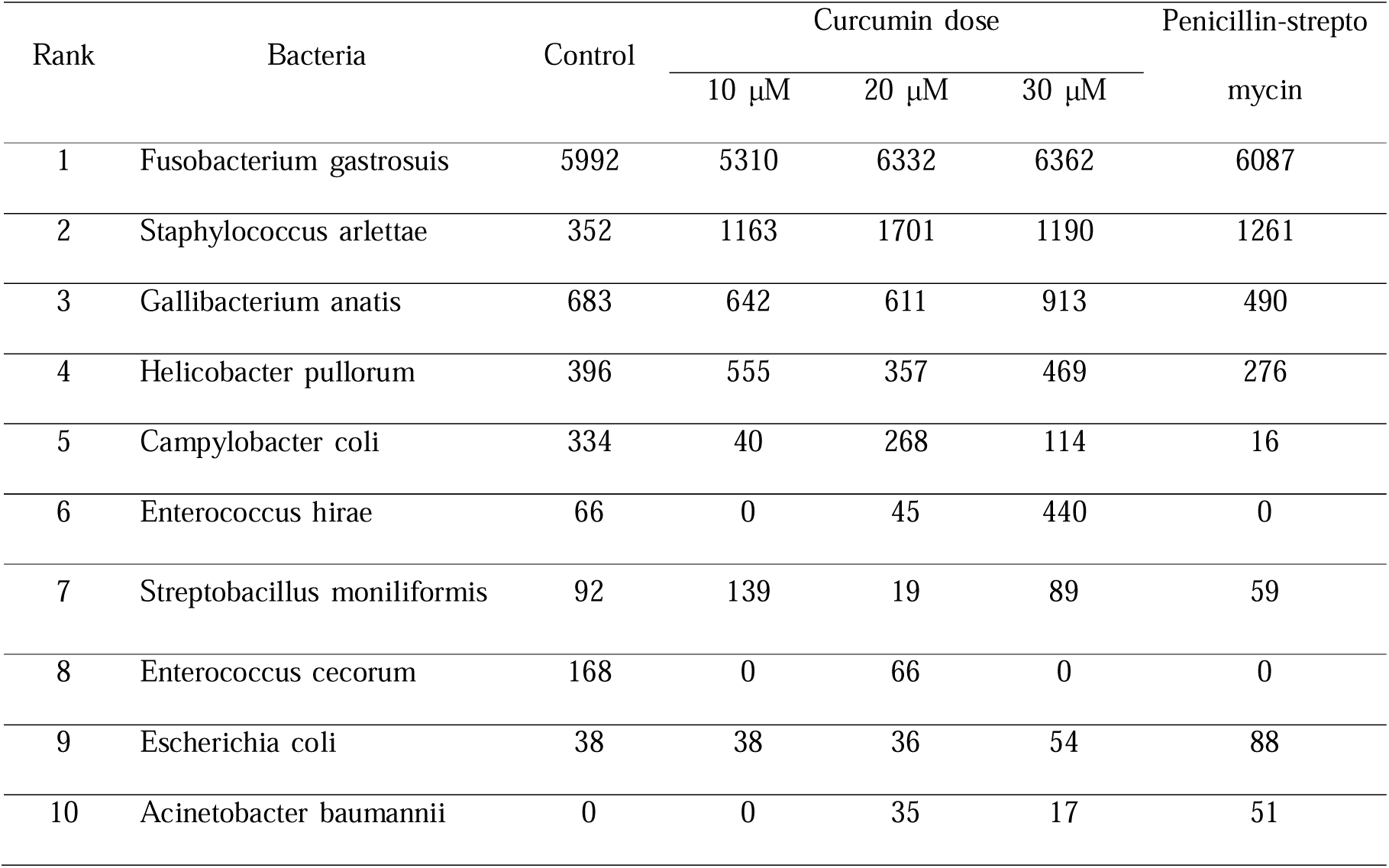
Read counts of pathogenic/pathobiont bacteria in rooster semen.

| Rank | Bacteria | Control | Curcumin dose |  |  | Penicillin-strepto<br>mycin |
| --- | --- | --- | --- | --- | --- | --- |
| | | | 10 μM | 20 μM | 30 μM | |
| 1 | *Fusobacterium gastrosuis* | 5992 | 5310 | 6332 | 6362 | 6087 |
| 2 | *Staphylococcus arlettae* | 352 | 1163 | 1701 | 1190 | 1261 |
| 3 | *Gallibacterium anatis* | 683 | 642 | 611 | 913 | 490 |
| 4 | *Helicobacter pullorum* | 396 | 555 | 357 | 469 | 276 |
| 5 | *Campylobacter coli* | 334 | 40 | 268 | 114 | 16 |
| 6 | *Enterococcus hirae* | 66 | 0 | 45 | 440 | 0 |
| 7 | *Streptobacillus moniliformis* | 92 | 139 | 19 | 89 | 59 |
| 8 | *Enterococcus cecorum* | 168 | 0 | 66 | 0 | 0 |
| 9 | *Escherichia coli* | 38 | 38 | 36 | 54 | 88 |
| 10 | *Acinetobacter baumannii* | 0 | 0 | 35 | 17 | 51 |

The group of commensal/beneficial-associated bacteria (Table 2) showed the dominance of several taxa that may play a role in fermentation and the balance of the microbial ecosystem. Ligilactobacillus salivarius was one of the most abundant bacteria, but its abundance decreased from the control to the 20 μM curcumin treatment and increased slightly at 30 μM curcumin. Ligilactobacillus aviarius and Lactobacillus crispatus also tended to decrease under curcumin treatment compared to the control. In contrast, Peptacetobacter hiranonis increased from the control, reaching its highest abundance at 20 μM curcumin, while Kineothrix sp. MB12-C1 also reached its highest value at 20 μM curcumin before decreasing at 30 μM curcumin. These patterns indicate that the response of commensal bacteria to curcumin is taxon-specific, with 20 μM curcumin appearing to provide relatively favorable conditions for certain fermentative bacteria, although not all commensal bacteria responded positively.

**Table 2.** Read counts of Nonpathogenic bacteria in rooster semen.

| Rank | Bacteria | control | Curcumin dose |  |  | Penicillin-strepto<br>mycin |
| --- | --- | --- | --- | --- | --- | --- |
| | | | 10 μM | 20 μM | 30 μM | |
| 1 | *Porphyromonas somerae* | 11587 | 12794 | 12355 | 11951 | 9332 |
| 2 | uncultured *Saccharofermentans* sp. | 7930 | 11129 | 9367 | 12475 | 11545 |
| 3 | *Kineothrix* sp. MB12-C1 | 7781 | 8129 | 9651 | 6942 | 7397 |
| 4 | *Ligilactobacillus salivarius* | 10628 | 8464 | 4924 | 5692 | 4227 |
| 5 | *Mageeibacillus indolicus* | 4828 | 5128 | 4313 | 4903 | 3473 |
| 6 | *Ligilactobacillus aviarius* | 2008 | 1057 | 1443 | 897 | 1753 |
| 7 | *Lactobacillus crispatus* | 1624 | 1048 | 1282 | 869 | 2088 |
| 8 | *Peptacetobacter hiranonis* | 810 | 1086 | 1551 | 1213 | 400 |
| 9 | *Ligilactobacillus agilis* | 1708 | 939 | 753 | 618 | 858 |
| 10 | *Olsenella* sp. An293 | 548 | 686 | 438 | 1125 | 873 |

### Alpha Diversity

The number of observed species indicates that the control group had the highest species diversity (1,210), while the curcumin and penicillin-streptomycin treatments reduced species diversity, with the lowest value observed in the 30 μM curcumin treatment (986). Estimates of species richness using the Chao1 index showed that the control group (1,449.71) had the highest value, while the lowest value was observed in the 30 μM curcumin group (1,225.04). Based on the ACE index, there were differences in estimates of microbial community richness, with the control group having the highest ACE index (1,498.11), while the lowest ACE value was found in the 30 μM curcumin group (1,286.53). Based on the Shannon and Simpson indices, all samples showed relatively uniform levels of microbial diversity and evenness, with values ranging from 4.07 to 4.21 and 0.96, respectively (Figure 6).

**Figure 6.**
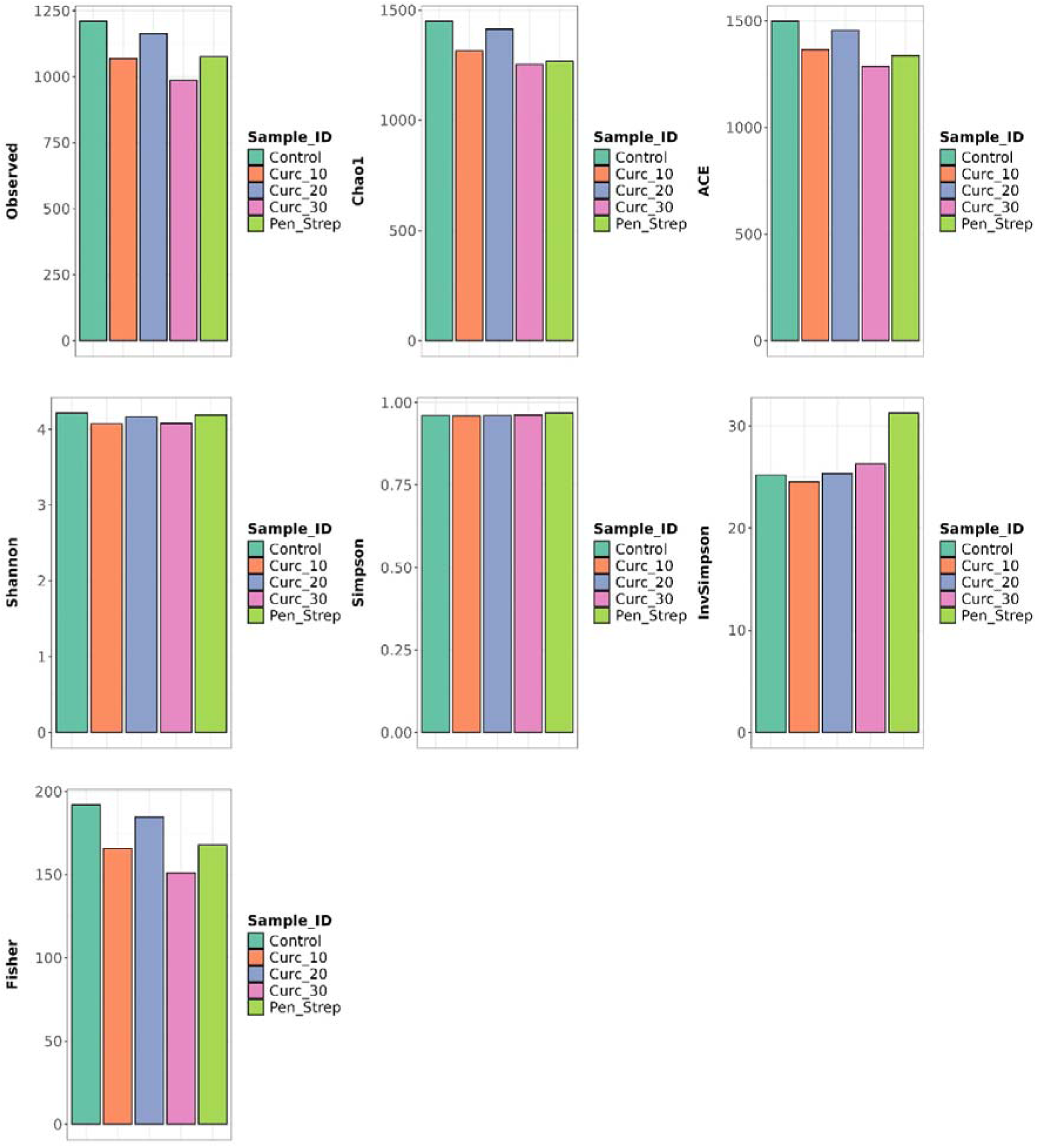
Results of alpha diversity analysis in rooster semen.

Based on the Inverse Simpson index graph, the penicillin-streptomycin group showed higher values than the other groups, indicating that the use of penicillin-streptomycin resulted in the most optimal level of microbial community evenness. The relatively high Fisher index values (>150) in all samples indicate that the samples were collected from environments or ecosystems rich in biodiversity.

### Beta Diversity

Beta diversity analysis using Principal Component Analysis (PCA) (Figure 7) shows that the PC1, PC2, and PC3 axes explain 30.37%, 28.26%, and 22.03% of the total biological diversity across areas, respectively. Based on the PC1 vs. PC2 ordination plot, the 10 μM curcumin and 20 μM curcumin samples are located close together in the same quadrant, indicating a high degree of similarity in species composition. In contrast, the 30 μM curcumin, control, and penicillin-streptomycin samples are separated from one another in PCA space, indicating a high level of biological diversity (beta diversity). Meanwhile, in the PC1 vs. PC3 plot, the control group is in the same quadrant as the 20 μM curcumin, while the 10 μM curcumin, 30 μM curcumin, and penicillin-streptomycin are separated from one another in PCA space. Overall, the distribution of the five sample points illustrates a pattern of ecosystem differentiation influenced by curcumin concentration and antibiotic use.

**Figure 7.**
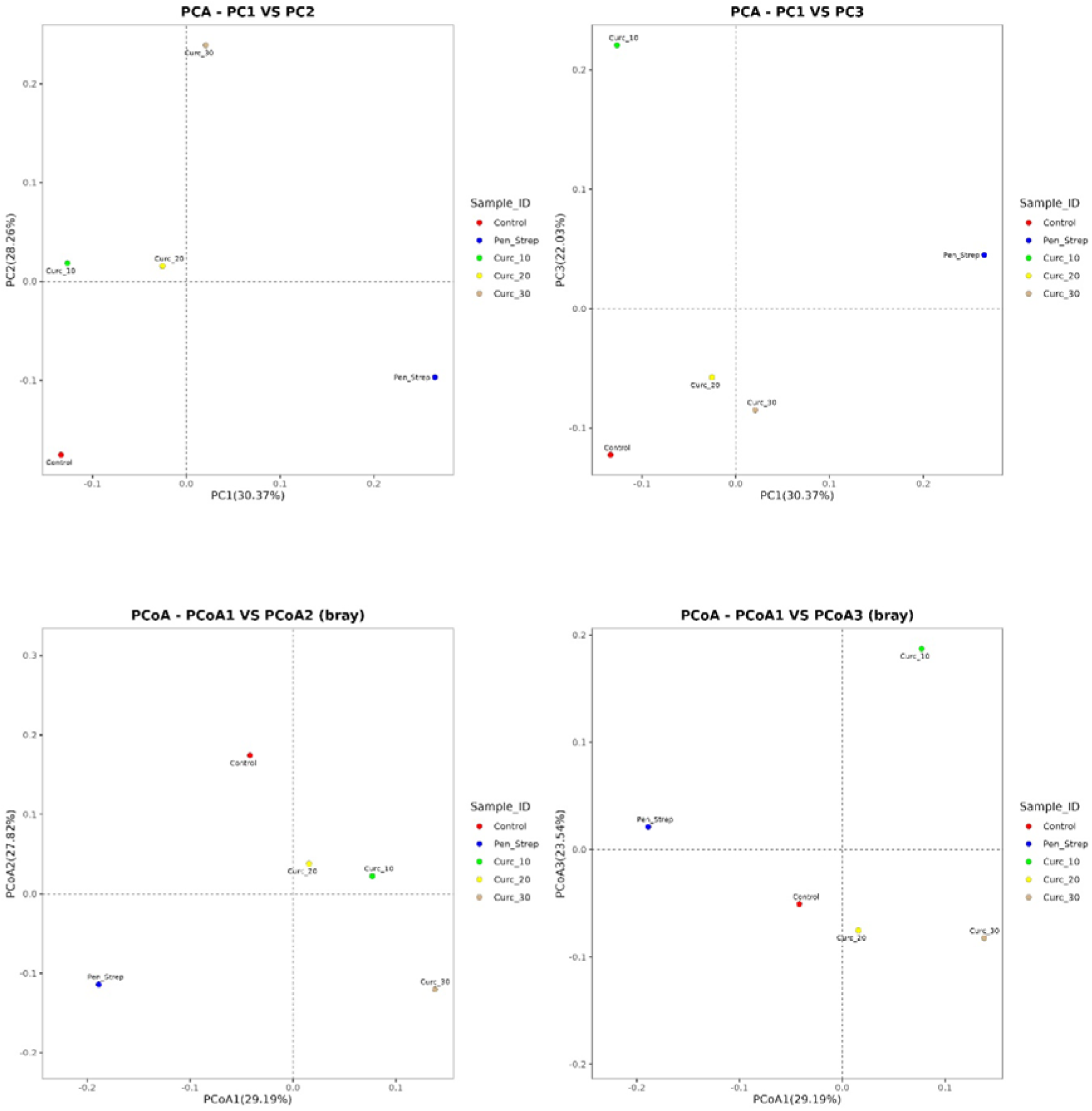
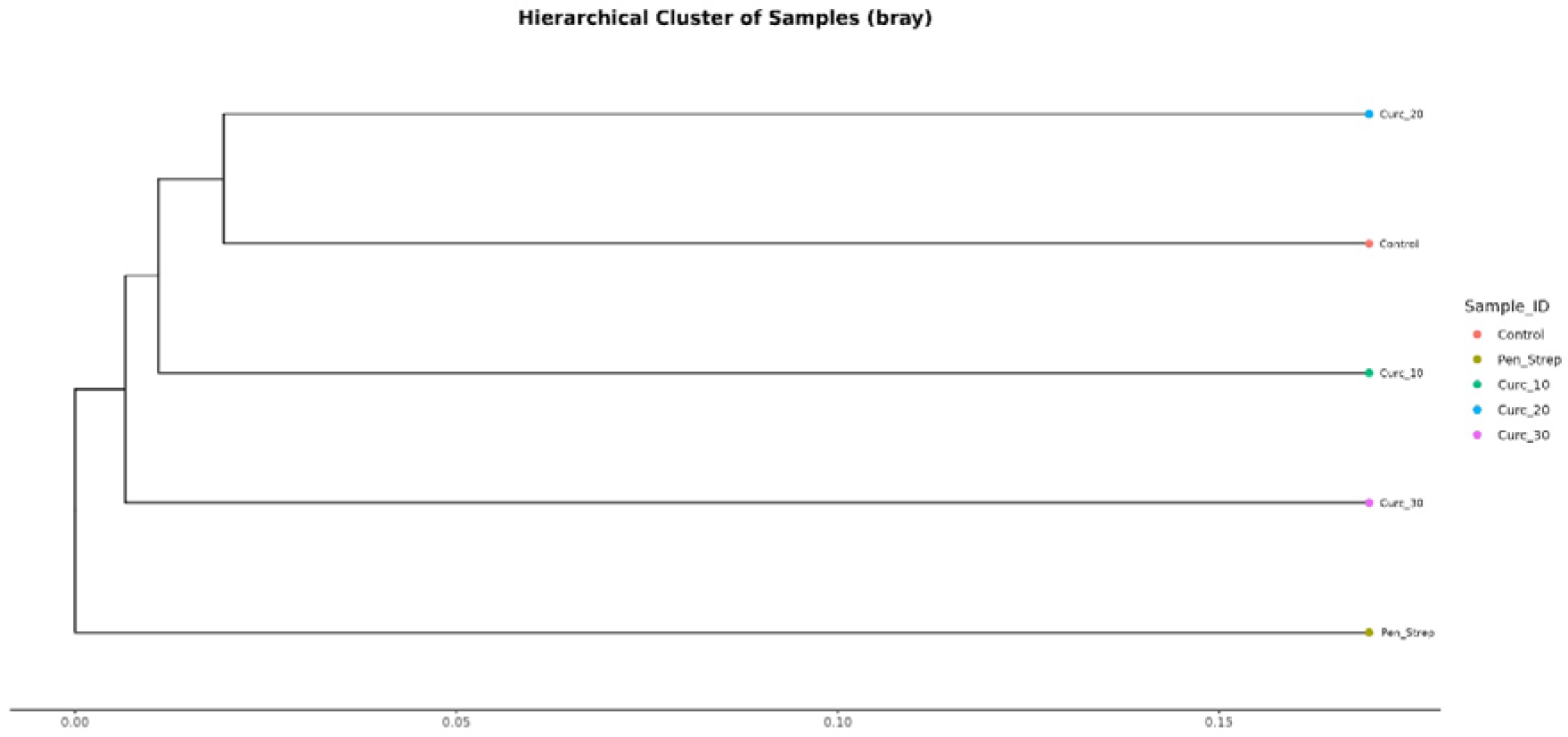
Figure 7. Results of beta diversity analysis in rooster semen. Top: Principal Component Analysis (PCA); Middle: Principal Coordinate Analysis (PCoA); Bottom: Unweighted Pair Group Method with Arithmetic Mean (UPGMA)-based hierarchical clustering analysis.

An analysis of beta diversity using Principal Coordinate Analysis (PCoA) based on the Bray-Curtis distance matrix (Figure 7) revealed distinct community structures among the five samples. The PCoA1, PCoA2, and PCoA3 axes cumulatively explain 29.19%, 27.82%, and 23.54% of the data variance, respectively. Based on the PCoA1 vs. PCoA2 ordination plot, the 10 μM and 20 μM curcumin samples are close to each other in the upper right quadrant, indicating a high degree of similarity in biological composition. However, the 20 μM curcumin sample exhibits a closer relationship with the 30 μM curcumin sample based on the secondary factor along the PCoA3 axis (23.54%), which significantly distinguishes the 20 μM and 30 μM curcumin samples from the 10 μM curcumin sample. Thus, the 20 μM curcumin sample occupies an intermediate position linking the characteristics of the 10 μM and 30 μM curcumin samples.

The results of the UPGMA-based hierarchical clustering analysis (Figure 7) show that the addition of penicillin-streptomycin caused the most significant change in the profile, as indicated by its separation as a major outlier from all other sample groups. In contrast, the samples treated with curcumin clustered closely with the control samples, with the 20 μM curcumin treatment showing the highest similarity to the control.

### Heatmap

Saccharofermentans sp. (which has not yet been cultured), Porphyromonas somerae, and Kineothrix sp. MB12-C1 accounted for approximately 10% of the total population in all study samples. Ligilactobacillus salivarius, at 10%, was found in the control sample and the 10 μM curcumin sample. Some species accounted for only 1% of the total population, such as Helococcus ovis, Erysipelothrix sp. HDW6B, and Holdemania massiliensis. The concentration of Qiania dongpingensis was very low (0.001%) in the 20 μM curcumin sample, whereas in the other samples, its concentration was 0.1% (Figure 8).

**Figure 8.**
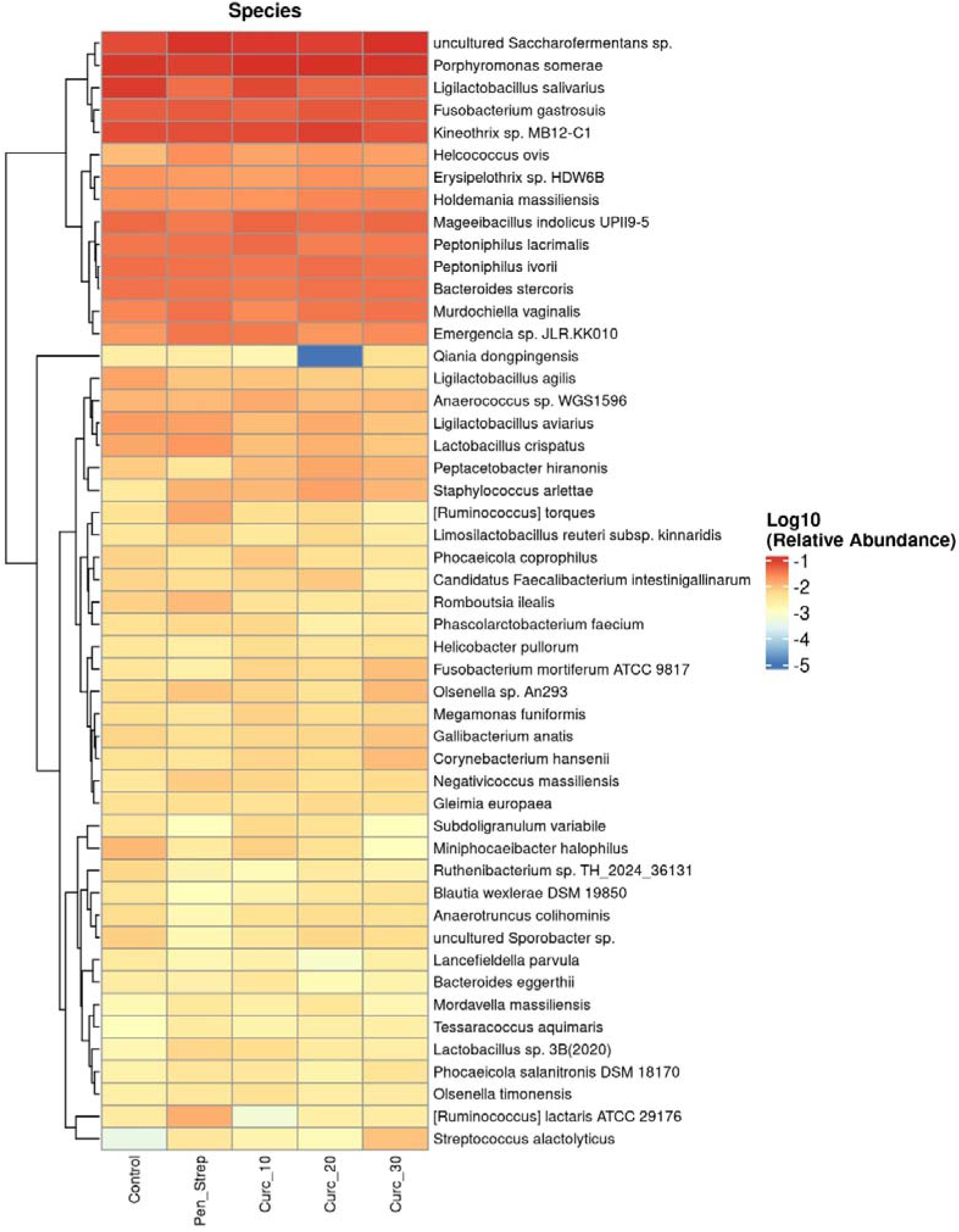
Heat map of bacteria in rooster semen.

## DISCUSSION

This study has provided new insights into the identification of microbiome community structure and taxonomic diversity in chicken semen diluted with curcumin and penicillin-streptomycin. To the best of our knowledge, this study is the first to characterize the microbiome profile of diluted chicken semen stored at 5 °C for 24 hours using a 16S full-length amplicon sequencing approach, which allows for taxonomic resolution down to the species level.

A two-way Venn diagram analysis showed that the administration of penicillin-streptomycin caused a decrease in natural taxonomic richness, with 441 OTUs from the control group no longer detected after penicillin-streptomycin treatment. This decrease reflects the bactericidal effect of penicillin-streptomycin on susceptible microbial groups. Streptomycin works by inhibiting bacterial protein synthesis [18], and penicillin functions by inhibiting the cross-linking of peptidoglycan within the bacterial cell wall [19]. Analysis of microbial community intersections using a 5-way Venn diagram showed that each treatment group exhibited the emergence of unique OTUs reflecting their respective shifts in microbial community profiles. The control group retained 117 unique OTUs, reflecting natural diversity; the use of penicillin-streptomycin resulted in 109 unique OTUs, indicating the occurrence of profound dysbiosis, in which resistant or opportunistic microbial groups dominate new ecological niches following the elimination of sensitive taxa [20]. The decrease in the number of unique OTUs with the use of 30 μM curcumin indicates that higher concentrations of curcumin act through a selective elimination mechanism that enhances certain beneficial taxa while suppressing the diversity of non-specific taxa. Curcumin alters the microbiome ecosystem, thereby promoting increased production of beneficial short-chain fatty acid-producing taxa and a decrease in the Firmicutes/Bacteroidetes ratio [21].

Overall, the Sankey analysis showed that curcumin treatment did not fundamentally alter the structure of the chicken semen microbial community at the phylum level, as Bacillota remained the dominant group and Bacteroidota remained the next most dominant group across all treatments. Changes were more evident in the distribution and relative abundance of taxa at the class through genus levels, including variations in Clostridia, Bacilli, Tissierellia, Lactobacillaceae–Ligilactobacillus, Porphyromonadaceae–Porphyromonas, and several other anaerobic groups. In the chicken semen microbiome in this study, Bacillota, Bacteroidota, and Fusobacteriota were the most abundant phyla. These results differ slightly from the report by Jiang et al. [4], which found that the phyla Actinobacteria, Bacteroidota, Firmicutes, and Proteobacteria were the most abundant in chicken semen in China. At the genus level, this study identified Ligilactobacillus, Porphyromonas, Peptoniphilus, Saccharofermentans, Kineothrix, and Fusobactorium as the most abundant. Meanwhile, Jiang et al. [4] reported that the most abundant bacterial genera in Chinese chicken semen were Lactobacillus, Anaerococcus, Actinomyces, Peptoniphilus, Phyllobacterium, and Porphyromonas. Tvrdá et al. [15] reported that the genera Escherichia, Staphylococcus, Citrobacter, and Enterococcus were identified as the most abundant in Lohmann Brown and Ross chickens. Meanwhile, Klebsiella and Achromobacter were detected in the semen of native Thai chickens [22]. Furthermore, Ďuračka et al. [6] found that the most dominant bacterial genus in Lohmann rooster semen was Enterococcus. This is understandable because variations in semen bacterial profiles depend on the rooster breed [22]. The presence of several commensal and saccharolytic taxa, such as Lactilactobacillus salivarius, Kineothrix sp. MB12-C1, uncultured Saccharofermentans sp., and Bacteroides stercoris, indicates active fermentative metabolic activity in seminal plasma. The dominance of these taxa suggests the possible presence of carbohydrate substrates in seminal plasma as well as the potential formation of metabolites such as lactic acid and short-chain fatty acids (SCFAs)—including propionate, acetate, and butyrate—which maintain microbiome balance by suppressing pathogenic bacteria. Kineothrix is known to produce butyric acid [23], Saccharofermentans produces lactic acid and acetate [24], Bacteroides produces propionate and acetate [25], and Ligilactobacillus produces lactic acid [26]. Ligilactobacillus salivarius was quite abundant in the control group and the 20 μM curcumin group. Ligilactobacillus salivarius is found in the gastrointestinal tract of chickens and acts as a probiotic; this bacterium produces salivarin, a bacteriocin with broad-spectrum antipathogenic effects that effectively inhibits pathogenic bacteria such as Salmonella [27], Campylobacter [28], and Escherichia coli [29]. It is known that Salmonella, Campylobacter, and Escherichia coli can reduce the motility of chicken spermatozoa [7]. Therefore, a high abundance of L. salivarius in seminal plasma is actually beneficial for spermatozoa.

In addition, the abundance of acidolytic taxa such as Peptoniphilus ivorii and Peptoniphilus lacrimalis reflects the utilization of protein-and amino acid-based substrates in chicken manure. Peptoniphilus uses peptone and amino acids as its primary energy sources, with butyric acid as the end product of metabolism [30]. The presence of Fusobacterium gastrosuis, Porphyromonas somerae, and Murdochiella vaginalis indicates the existence of an ecological niche associated with the mucosa or a strictly anaerobic environment. Although some members of these genera have the potential to act as opportunistic pathogens under conditions of dysbiosis, their presence in these samples may reflect part of the natural microflora diversity that has adapted to the physicochemical conditions of chicken semen.

The use of curcumin at various concentrations tended not to cause changes in the relative abundance of Porphyromonas somerae, presumably because the curcumin concentrations were still low enough to inhibit the growth of this bacterium. However, a slight decrease in population was clearly observed in the penicillin-streptomycin group. Although the effects of penicillin on Porphyromonas somerae have not been previously studied, Navarrete-López et al. [31] confirmed that Porphyromonas may be beneficial because it is present in the semen of fertile horses; this suggests that this bacterium could serve as a good indicator of optimal semen quality. Therefore, further studies are needed regarding the effects of this bacterium’s presence on chicken sperm quality.

Saccharofermentans sp. was found in low percentages in the control group and showed an increase in the curcumin and penicillin-streptomycin groups in this study. Saccharofermentans is a non-pathogenic, Gram-positive, non-motile, obligate anaerobic bacterium that does not exhibit microaerophilic or aerobic growth; it grows at neutral pH and is a chemoorganotroph capable of fermenting glucose into acetate, lactate, and fumarate [33]. Therefore, the presence of this bacterium is unlikely to have a negative impact on the viability of chicken sperm.

Overall, curcumin treatment did not show a consistent pattern of inhibition against all pathogenic/pathobiont bacteria, but produced a specific response to each taxon and was not linear with increasing concentration. Previous studies have shown that Bifidobacterium, Salmonella, Campylobacter, E. coli, and Clostridium reduce sperm motility in broiler chickens [7]. The use of penicillin-streptomycin slightly increased Escherichia in this study, which may be due to resistance, as it has been reported that Escherichia coli isolated from chickens exhibits increased resistance to streptomycin [34] and penicillin [35].

The metagenomic alpha diversity profile of chicken semen reveals quite interesting dynamics in response to curcumin and penicillin-streptomycin treatments. Species richness indices such as Observed, Chao1, ACE, and Fisher consistently yielded the highest values in the control group, then decreased as curcumin concentration increased and in the penicillin-streptomycin group. This decrease in microbial richness indicates that curcumin possesses broad-spectrum antimicrobial activity capable of suppressing the abundance of certain sensitive microbial taxa within the ecosystem [12], as does penicillin-streptomycin [36]. This dose-dependent reduction in abundance is consistent with curcumin’s ability to damage cell membranes and disrupt bacterial cell division [37].

Interestingly, despite a decline in species richness, the Shannon and Simpson diversity indices remained relatively constant across all treatment groups. This suggests that although the curcumin and penicillin-streptomycin treatments eliminated or suppressed rare microbes, the underlying structure and evenness of the core microbiome community in chicken semen remained balanced. On the other hand, the InvSimpson index showed the highest value in the penicillin-streptomycin group, indicating a decrease in the dominance of certain taxa and an increase in the evenness of resistant/tolerant microbial groups following antibiotic exposure. Antibiotic-induced changes in microbial composition include reduced microbial diversity, alterations in the functional attributes of the microbiota, and the formation and selection of antibiotic-resistant strains [38]. Overall, curcumin exhibits a suppressive effect on microbial richness comparable to that of antibiotics, which explains its potential as a selective phytobiotic agent in modulating the chicken semen microbiota without drastically disrupting the ecological balance. Thus, curcumin can serve as a beneficial natural alternative to antibiotics, thereby contributing to efforts to address antibiotic resistance [13].

Beta-diversity ordination analysis confirmed that the penicillin-streptomycin treatment caused the most drastic shift in microbial community structure, resulting in a group that was far more isolated from the others. In PCoA, this separation reflects a loss of diversity as well as changes in taxon dominance, whereas in PCA, the separation along the PC1 axis explains the occurrence of extreme changes in the quantitative abundance of bacteria on a massive scale. This effect is consistent with the phenomenon of antibiotic-induced dysbiosis, in which antibiotic exposure severely disrupts this ecosystem by reducing microbial diversity and depleting beneficial commensal bacteria [39]. In contrast, the curcumin group demonstrated a gradual, dose-dependent modulation mechanism. In both the PCoA1 vs. PCoA2 and PCA1 vs. PC3 plots, the 10 μM, 20 μM, and 30 μM curcumin samples moved along a gradient trajectory away from the control. This pattern indicates that curcumin does not act as a mass killer like synthetic antibiotics, but rather as a selective modulator that gradually restructures the abundance of certain taxa—such as suppressing opportunistic pathogenic bacteria while supporting the growth of beneficial bacteria [40]. Hierarchical clustering analysis using the UPGMA method showed that the penicillin-streptomycin group formed a separate branch at the base of the tree (outgroup), confirming that antibiotic exposure triggered the most drastic changes in taxonomic composition compared to the curcumin intervention group. Meanwhile, the control and 20 μM curcumin treatments clustered within a single major subcluster, indicating that the underlying structure of the microbial community still shared characteristics similar to those of the control condition. The merging of the 10 μM and 30 μM curcumin treatments at the next branch confirms a dynamic yet controlled pattern of change. Unlike penicillin-streptomycin, which triggered massive and extreme branching, curcumin administration gradually modulated the microbiome profile without massively disrupting the underlying structure of the microflora ecosystem.

Implicitly, these findings demonstrate that curcumin has high potential as a candidate for modulating the natural microbial community—one that is safer than penicillin-streptomycin—because it can steer the structure of the microflora toward a new profile without severely disrupting the stability of the ecosystem.

## AUTHORS’ CONTRIBUTION

Conceptualization: Khaeruddin, Hermawansyah

Data curation: Kasri

Formal analysis: Khaeruddin, Kasri

Methodology: Hermawansyah, Junaedi

Validation: Khaeruddin, Junaedi

Investigation: Hermawansyah

Writing - original draft: Syamsuryadi B, Kasri, Khaeruddin

Writing - review & editing: Khaeruddin, Hermawansyah, Junaedi, Syamsuryadi B, Kasri

## ACKNOWLEDGMENTS

The authors would like to thank the Directorate General of Research and Development, Ministry of Higher Education, Science, and Technology, for the financial support provided through the Fiscal Year 2026 Fundamental Research Grant Program. Thanks are also extended to PT Genetik Science Indonesia for hosting the metagenomic analysis.

## SUPPLEMENTARY MATERIAL

Not applicable.

## DATA AVAILABILITY

Upon reasonable request, the datasets of this study can be available from the corresponding author.

## ETHICS APPROVAL

This study has received ethical approval from the Animal and Research Ethics Committee at Hasanuddin University, as evidenced by Decision Letter No. 00246/UN4.1.30.1.1.4/DI.05.01/2026.

## DECLARATION OF GENERATIVE AI

During the preparation of this work, ChatGPT was used for the purpose of language editing and copyediting. After using this tool, the manuscript was reviewed and edited as needed and authors will assume full responsibility for the publication.

## REFERENCES

1. Mohan J, Kolluri G, Srivastava V, Tyagi JS, Tiwari AK. Bacterial contamination of poultry semen, its dilution and storage. Worlds Poult Sci J 2023;79(3):593–617. 10.1080/00439339.2023.2225793

2. Althouse GC. Sanitary procedures for the production of extended semen. Reprod Domest Anim 2008;43:374–8. 10.1111/j.1439-0531.2008.01187.x

3. Maung EE, Sushadi PS, Asano A. Polymyxin B neutralizes endotoxin and improves the quality of chicken semen during liquid storage. Theriogenology. 2023;198:107–13. 10.1016/j.theriogenology.2022.12.027

4. Jiang X, Zhang B, Gou Q, Cai R, Sun C, Li J, Yang N, Wen C. Variations in seminal microbiota and their functional implications in chickens adapted to high-altitude environments. Poult Sci 2024;103(8):103932. 10.1016/j.psj.2024.103932

5. Monteiro C, Marques PI, Cavadas B, Damião I, Almeida V, Barros N, Barros A, Carvalho F, Gomes S, Seixas S. Characterization of microbiota in male infertility cases uncovers differences in seminal hyperviscosity and oligoasthenoteratozoospermia possibly correlated with increased prevalence of infectious bacteria. Am J Reprod Immunol 2018;79(6):e12838. 10.1111/aji.12838

6. Ďuračka M, Petrovičová M, Benko F, Kováčik A, Lukáč N, Kačániová M, Tvrdá E. Lohmann Brown Rooster Semen: intrinsic bacteria and their impact on sperm progressive motility and seminal biochemical parameters—a preliminary study. Stresses. 2023;3(2):424–33. 10.3390/stresses3020031

7. Haines MD, Parker HM, McDaniel CD, Kiess AS. Impact of 6 different intestinal bacteria on broiler breeder sperm motility in vitro. Poult Sci. 2013;92(8):2174–81. 10.3382/ps.2013-03109

8. Tvrdá E, Petrovičová M, Benko F, Ďuračka M, Galovičová L, Slanina T, Kačániová M. Curcumin attenuates damage to rooster spermatozoa exposed to selected uropathogens. Pharmaceutics. 2022;15(1):65. 10.3390/pharmaceutics15010065

9. Khaeruddin K, Ciptadi G, Yusuf M, Udrayana SB, Iswati S, Wahjuningsih S. Effectiveness of butylated hydroxytoluene in maintaining the quality of Gaga chicken sperm in liquid storage for 72 hours. Adv Anim Vet Sci 2024;12(2):371–80. 10.17582/journal.aavs/2024/12.2.371.380

10. Morrell JM, Wallgren M. Alternatives to antibiotics in semen extenders: A review. Pathogens 2014;3(4):934–46. 10.3390/pathogens3040934

11. Lobanovska M, Pilla G. Penicillin’s discovery and antibiotic resistance: lessons for the future?. Yale J Biol Med 2017;90(1):135.

12. Cojkic A, Hansson I, Johannisson A, Morrell JM. Effect of some plant-based substances on microbial content and sperm quality parameters of bull semen. Int J Mol Sci 2023;24(4):3435. 10.3390/ijms24043435

13. Aderemi FA, Alabi OM. Turmeric (Curcuma longa): an alternative to antibiotics in poultry nutrition. Transl Anim Sci 2023;7(1):txad133. 10.1093/tas/txad133

14. Gil L, González N, Horndler L, Luño V. Dose-dependent effects of curcumin on bacterial growth and sperm quality during refrigerated storage of equine epididymal sperm. Front Vet Sci 2026;13:1739360. 10.3389/fvets.2026.1739360

15. Tvrdá E, Petrovičová M, Benko F, Ďuračka M, Kováč J, Slanina T, Galovičová L, Žiarovská J, Kačániová M. Seminal bacterioflora of two rooster lines: Characterization, antibiotic resistance patterns and possible impact on semen quality. Antibiotics 2023;12(2):336. 10.3390/antibiotics12020336

16. Kucera AC, Heidinger BJ. Avian semen collection by cloacal massage and isolation of DNA from sperm. J Vis Exp 2018;132:55324. 10.3791/55324

17. Chen YL, Lee CC, Lin YL, Yin KM, Ho CL, Liu T. Obtaining long 16S rDNA sequences using multiple primers and its application on dioxin-containing samples. BMC Bioinformatics 2015;16(Suppl 18):S13. 10.1186/1471-2105-16-S18-S13

18. He C, Eggelbusch M, Huijts JY, Shi A, de Wit GJ, Offringa C, Jaspers RT, Wüst RC. The commonly used antibiotic streptomycin reduces protein synthesis and differentiation in cultured C2C12 myotubes. Physiol Rep 2025;13(12):e70353. 10.14814/phy2.70353

19. Fisher JF, Mobashery S. Constructing and deconstructing the bacterial cell wall. Protein Sci 2020;29(3):629–46. 10.1002/pro.3737

20. Fukuyama J, Rumker L, Sankaran K, Jeganathan P, Dethlefsen L, Relman DA, Holmes SP. Multidomain analyses of a longitudinal human microbiome intestinal cleanout perturbation experiment. PLoS Comput Biol 2017;13(8):e1005706. 10.1371/journal.pcbi.1005706

21. Konaktchieva M, Stojchevski R, Hadzi-Petrushev N, Gagov H, Konakchieva R, Mitrokhin V, Kungulovski G, Mladenov M, Avtanski D. Curcumin and tetrahydrocurcumin as multi-organ modulators of the adipose tissue–gut–liver axis: mechanistic insights, therapeutic potential, and translational challenges. Pharmaceuticals 2025;18(12):1791. 10.3390/ph18121791

22. Authaida S, Boonkum W, Duangjinda M, Chankitisakul V. Comparative variation and associations among seminal microbiota, oxidative status, and semen quality in different rooster types. Animals 2026;16(9):1380. 10.3390/ani16091380

23. Haas KN, Blanchard JL. Kineothrix alysoides, gen. nov., sp. nov., a saccharolytic butyrate-producer within the family Lachnospiraceae. Int J Syst Evol Microbiol 2017;67(2):402–10. 10.1099/ijsem.0.001643

24. Monteiro HF, Lelis AL, Fan P, Agustinho BC, Lobo RR, Arce-Cordero JA, Dai X, Jeong KC, Faciola AP. Effects of lactic acid-producing bacteria as direct-fed microbials on the ruminal microbiome. J Dairy Sci 2022;105(3):2242–55. 10.3168/jds.2021-21025

25. Louis P, Flint HJ. Formation of propionate and butyrate by the human colonic microbiota. Environ Microbiol 2017;19(1):29–41. 10.1111/1462-2920.13589

26. Alba C, Arroyo R, Fernández L, Narbad A, Rodríguez JM. Characterization of a Ligilactobacillus salivarius strain isolated from a cheese seal which was last used in 1936. Foods. 2024 13(13):2005. 10.3390/foods13132005

27. Maniee SA, Mahmoodian S, Zamani Amirzakaria J, Meimandipour A, Shariati V, Tavakol E. Whole-genome analysis of Ligilactobacillus salivarius L33, a potential probiotic strain isolated from chicken gastrointestinal tract. Microbiol Spectr 2025;13(10):e01591–24. 10.1128/spectrum.01591-24

28. Stern NJ, Svetoch EA, Eruslanov BV, Perelygin VV, Mitsevich EV, Mitsevich IP, Pokhilenko VD, Levchuk VP, Svetoch OE, Seal BS. Isolation of a Lactobacillus salivarius strain and purification of its bacteriocin, which is inhibitory to Campylobacter jejuni in the chicken gastrointestinal system. Antimicrob Agents Chemother 2006;50(9):3111–6. 10.1128/aac.00259-06

29. Wang J, Ishfaq M, Guo Y, Chen C, Li J. Assessment of probiotic properties of Lactobacillus salivarius isolated from chickens as feed additives. Front Vet Sci 2020;7:415. 10.3389/fvets.2020.00415

30. Verma R, Morrad S, Wirtz JJ. Peptoniphilus asaccharolyticus-associated septic arthritis and osteomyelitis in a woman with osteoarthritis and diabetes mellitus. BMJ Case Rep 2017;2017:bcr-2017 219969. 10.1136/bcr-2017-219969

31. Navarrete-López P, Asselstine V, Maroto M, Lombó M, Cánovas Á, Gutiérrez-Adán A. RNA sequencing of sperm from healthy cattle and horses reveals the presence of a large bacterial population. Curr Issues Mol Biol 2024;46(9):10430–43. 10.3390/cimb46090620

32. Cojkic A, Hansson I, Johannisson A, Morrell JM. Effect of some plant-based substances on microbial content and sperm quality parameters of bull semen. Int J Mol Sci 2023;24(4):3435. 10.3390/ijms24043435

33. Chen S, Niu L, Zhang Y. Saccharofermentans acetigenes gen. nov., sp. nov., an anaerobic bacterium isolated from sludge treating brewery wastewater. Int J Syst Evol Microbiol 2010; 60(12):2735–8. 10.1099/ijs.0.017590-0

34. Besung IN, Sudipa PH, Suarjana IG, Suwiti N. Antibiotic resistance pattern of Escherichia coli isolated from layer chicken in Bali-Indonesia. J World’s Poult Res 2024;14(4):361–368. 10.36380/jwpr.2024.37

35. Akond M, Hassan SMR, Alam S, Shirin M. Antibiotic resistance of Escherichia coli isolated from poultry and poultry environment of Bangladesh. Am J Environ Sci 2009;5(1):47–52. 10.3844/ajessp.2009.47.52

36. Oplinger RW, Wagner EJ. Use of penicillin and streptomycin to reduce spread of bacterial coldwater disease I: antibiotics in sperm extenders. J Aquat Anim Health 2015;27(1):25–31. 10.1080/08997659.2014.966211

37. Dai C, Lin J, Li H, Shen Z, Wang Y, Velkov T, Shen J. The natural product curcumin as an antibacterial agent: Current achievements and problems. Antioxidants 2022;11(3):459. 10.3390/antiox11030459

38. Patangia DV, Anthony Ryan C, Dempsey E, Paul Ross R, Stanton C. Impact of antibiotics on the human microbiome and consequences for host health. Microbiologyopen 2022;11(1):e1260. 10.1002/mbo3.1260

39. Niculescu AG, Iacob CM, Brătilă E, Tocariu R, Coroleucă CA, Corcionivoschi N, Vrancianu CO, Popescu DL, Popa GL, Popa MI, Cristian RE. Antibiotic-driven gut microbiome dysbiosis: resistome dynamics, metabolic disruption, and paths to restoration. Antibiotics 2026;15(7):688. 10.3390/antibiotics15070688

40. Servida S, Piontini A, Gori F, Tomaino L, Moroncini G, De Gennaro Colonna V, La Vecchia C, Vigna L. Curcumin and gut microbiota: A narrative overview with focus on glycemic control. Int J Mol Sci 2024;25(14):7710. 10.3390/ijms25147710

